# Protein-Cadmium Interactions in Crowded Biomolecular Environments Probed by In-cell and Lysate NMR Spectroscopy

**DOI:** 10.1101/2023.11.03.565546

**Authors:** Sachin S. Katti, Tatyana I. Igumenova

## Abstract

One of the mechanisms by which toxic metal ions interfere with cellular functions is ionic mimicry, where they bind to protein sites in lieu of native metals Ca^2+^ and Zn^2+^. The influence of crowded intracellular environments on these interactions is not well understood. Here, we demonstrate the application of *in-cell* and lysate NMR spectroscopy to obtain atomic-level information on how a potent environmental toxin cadmium interacts with its protein targets. The experiments, conducted in intact *E. coli* cells and their lysates, revealed that Cd^2+^ can profoundly affect the quinary interactions of its protein partners, and can replace Zn^2+^ in both labile and non-labile protein structural sites without significant perturbation of the membrane binding function. Surprisingly, in crowded molecular environments Cd^2+^ can effectively target not only all-sulfur and mixed sulfur/nitrogen but also all-oxygen coordination sites. The sulfur-rich coordination environments show significant promise for bioremedial applications, as demonstrated by the ability of the designed protein scaffold α_3_DIV to sequester intracellular cadmium. Our data suggests that *in-cell* NMR spectroscopy is a powerful tool for probing interactions of toxic metal ions with their potential protein targets, and for the assessment of potency of sequestering agents.

## Introduction

Nutritive, or native, metal ions are essential for life-sustaining biological processes.^[1]^ Metal ion-binding events affect the structural and functional properties of biological macromolecules, thereby regulating catalysis,^[2]^ signal transduction,^[3]^ and energy generation.^[4]^ The selectivity of proteinic metal-ion interaction sites is defined by the structure of the first coordination shell and Lewis acidity of the ligands.^[5]^ Xenobiotic metals can subvert these selectivity barriers through ionic mimicry, where they directly compete with native metal ions for the binding sites in biological macromolecules. These non-native interactions interfere with vital cellular processes whose dysregulation poses significant threat to human health. Among notable examples, chronic exposure to potent environmental toxin Cd^2+^ is associated with neurological, respiratory, renal, and reproductive disorders.^[6]^

The structural analysis of Cd^2+^-bound proteins in the Protein Data Bank (PDB) reports that Cd^2+^ is found in a variety of coordination environments comprising O, S, and N ligands, and can adopt coordination numbers ranging from 4 to 8.^[7]^ Consistent with this finding, several *in vitro* studies have demonstrated the ability of Cd^2+^ to effectively displace Zn^2+^ and Ca^2+^ from the thiol- and oxygen-rich sites of proteins and protein domains.^[8]^ However, Cd^2+^ interactions in cellular environments can be affected by several factors such as influx-efflux homeostasis, accessibility of target sites, nutritive metal-ion pools, and intracellular chelators.^[9]^ To establish biological relevance, specificity and affinity of Cd^2+^ towards putative protein targets need to be examined in environments reflecting these conditions.

In crowded environments, detection of protein spectra and binding events requires advanced isotope enrichment schemes and rapid multidimensional data acquisition methods.^[10]^ Notable demonstrations of *in-cell* NMR spectroscopy applied to the studies of protein-metal ion interactions include Zn^2+^ binding to Cu/Zn superoxide dismutase 1 in HEK293T cells,^[11]^ and characterization of Ca^2+^-induced conformational rearrangement of calmodulin in *Xenopus laevis* oocytes.^[12]^ We have previously demonstrated that in buffered solutions, amide ^1^H and ^15^N NMR chemical shifts in proteins are exquisitely sensitive to the changes in electrostatic environment imparted by Cd^2+^ binding, leading to the unambiguous identification of Cd^2+^ binding sites.^[8b-e]^

The design of *in-cell* NMR studies to probe protein-Cd^2+^ interactions requires a live cell system resistant to heavy metals. *E. coli* cells exhibit substantial tolerance towards heavy metals like Cd^2+^, which is attributed to their evolved and adaptable resistance mechanisms.^[13]^ In this work, we present an approach to investigate protein-Cd^2+^ interactions in crowded biomolecular environments provided by *E. coli* cells and lysates (**Figure 1A**). In this approach, Cd^2+^ (or any other divalent metal ion) is added directly to the *E. coli* cell culture transformed with a plasmid that carries the target protein gene under the control of an inducible promoter. Addition of metal ions takes place after cells are transferred to the isotopically enriched growth medium and is timed to the induction of protein expression. Using 2D ^1^H-^15^N chemical shift correlation NMR spectroscopy, the effect of metal ions on the target protein is monitored directly in intact cells and lysates. Here, we apply this approach to three metal-binding protein systems to evaluate their susceptibility to Cd^2+^ interactions, establish the effect of Cd^2+^ on the quinary structure, and assess their potential as sequestering agents and sensors of cellular Cd^2+^ uptake.

**Figure 1.**
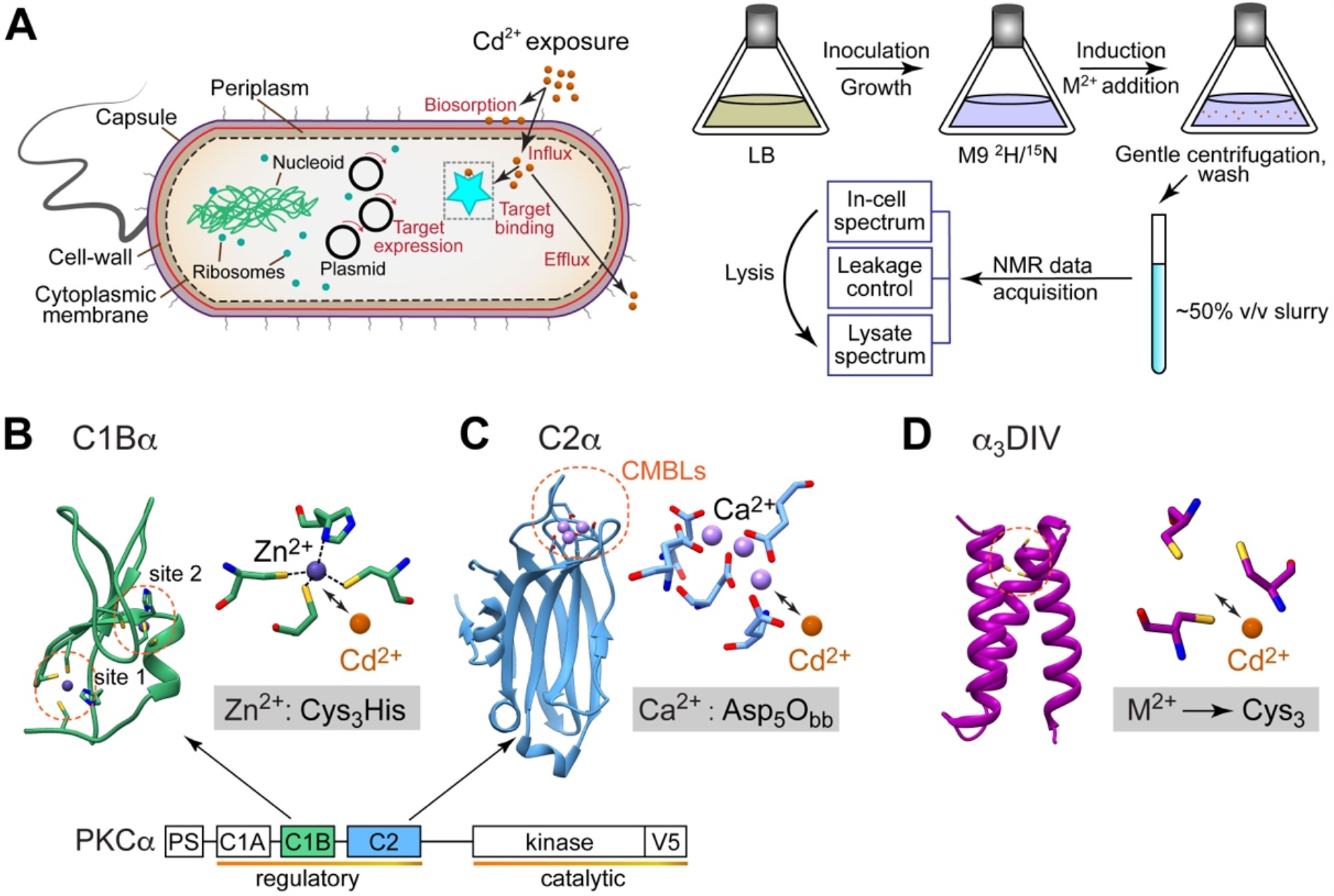
Overview of the experimental approach and protein systems for Cd^2+^ studies. (A) Experimental design for the detection of protein-Cd^2+^ interactions in intact E. coli cells and their lysates. NMR samples contain a suspension of intact E. coli cells that express target proteins and are grown in the isotopically-enriched medium with M^2+^ (M=Cd, Zn, Ca) supplementation. (B-D) Three-dimensional structures and metal-ion coordination sites of (B) C1Bα (PDB:2ELI), (C) C2α (PDB:3GPE), and (D) α3DIV (PDB:2MTQ). The modular diagram of PKCα shows regulatory and catalytic regions, with C1Bα and C2α domains color-coded green and blue, respectively.

## Results and discussion

### Choice of Protein Systems

The three protein systems chosen for this study present different metal-ion coordination environments to probe Cd^2+^ interactions, and have different folds, ligand preferences, and electrostatic surface properties. The first two proteins are independently folded peripheral membrane-binding modules of the conventional α isoform of protein kinase C (PKCα): the second cysteine-rich conserved homology-1 domain (C1Bα) and the conserved homology-2 domain (C2α) (**Figure 1B-C**). PKCα is a Ca^2+^-dependent Ser/Thr protein kinase of the AGC family. This enzyme is a regulatory node of the GPCR-mediated phosphoinositide signaling pathway and a proposed molecular target of Cd^2+^.^[14]^ Activation of PKCα requires membrane recruitment of its regulatory region comprising C1 and C2 domains, both of which contain metal ion binding sites that could be targeted by Cd^2+^ in the cellular environment. In C1Bα, these are represented by two Cys_3_His motifs, each coordinating a structural Zn^2+^ ion (**Figure 1B**).^[15]^ In C2α, it is the aspartate-rich Ca^2+^- and membrane-binding loops (CMBLs) that are of interest (**Figure 1C**).^[16]^ The third protein system is an engineered heavy metal-binding peptide, α_3_DIV, designed by Pecoraro’s laboratory (**Figure 1D**).^[17]^ This synthetic helical scaffold contains a pH-dependent, self-organizing Cys_3_ motif that mimics thiol-rich sites of metalloproteins, such as metallothionines.^[18]^ We and others have demonstrated all three proteins bind Cd^2+^ with high-affinity in aqueous buffer solutions.^[8b, 17]^

### NMR Detection of Target Proteins in Intact Cells

The first step was to establish the feasibility of detecting Cd^2+^-free C1Bα, C2α, and α_3_DIV in intact cells and lysates using 2D [^1^H,^15^N] NMR spectroscopy. The general sample preparation workflow involved recombinant expression of target proteins in *E. coli*, with cell cultures grown on the M9 minimal medium supplemented with [^15^N-enriched] NH_4_Cl. The growth medium was prepared in 100% D_2_O to achieve protein deuteration and thereby reduce NMR peak linewidths by attenuating ^1^H-mediated cross-relaxation pathways.

The 2D [^15^N-^1^H] SOFAST-HMQC NMR spectra collected on the 50% v/v slurry of intact cells show significant broadening of the ^15^N-^1^H_N_ resonances (**Figures 2A-C**). This is consistent with the restricted molecular tumbling in a high-viscosity cellular milieu, caused by proteins engaging in transient, low-affinity “quinary” interactions with endogenous macromolecules.^[19]^ Control NMR spectra on supernatants of cell suspensions were recorded after each *in-cell* experiment to ensure cell integrity and no protein leakage (representative spectrum is shown in **Figure S1A**).

**Figure 2.**
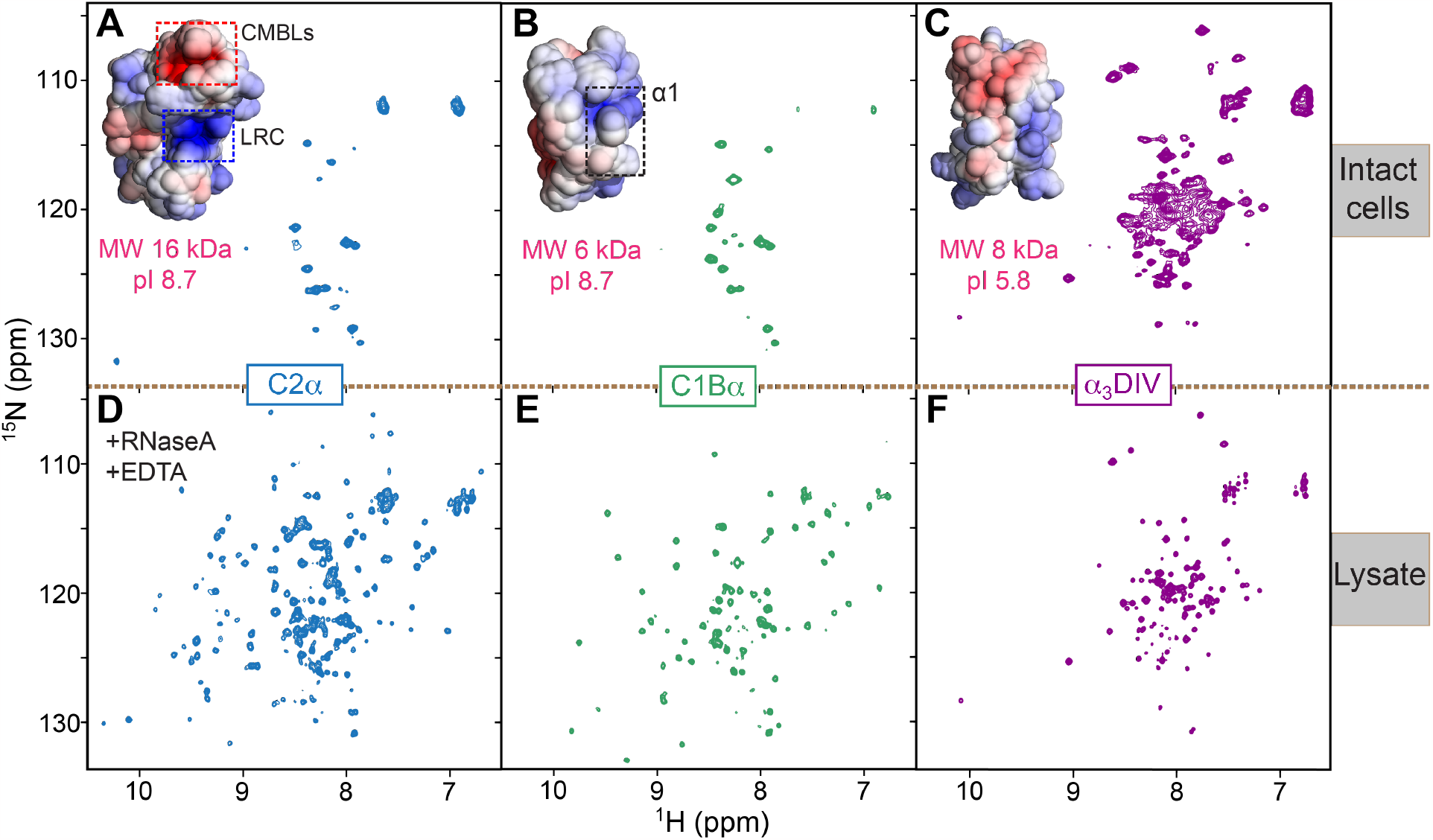
2D NMR detection of target proteins in intact E. coli cells and lysates. The in-cell/lysate [^15^N-^1^H] SOFAST-HMQC spectral pairs are shown for C2α (A,D), C1Bα (B,E), and α_3_DIV (C,F). The protein surfaces are color-coded according to the APBS-calculated electrostatic potential (± 5kT/e, insets of A-C). CMBL and LRC regions of C2α, and the α1 helix of C1Bα are highlighted with dashed boxes in (A) and (B), respectively. The amide chemical shift pattern in the in-cell α_3_DIV spectrum (C) is characteristic of an α-helical fold. Significant broadening of the amide resonances is observed for C2α and C1Bα in intact cells. All lysate spectra (D-F) report on fully folded proteins.

We find that the *in-cell* α_3_DIV spectrum contains many amide cross-peaks whose overall chemical shift dispersion and pattern report on a well-folded helical structure (**Figure 2C**). In contrast, the C2α and C1Bα data contain only a small subset of protein and metabolite cross-peaks in a narrow spectral range centered around ∼8.3 ppm (**Figure 2A, B**). To evaluate if C2α and C1Bα are properly folded, we conducted NMR experiments on the clarified lysates generated from the *in-cell* NMR samples. Persistent molecular crowding in lysates was verified by SDS-PAGE (**Figure S1B**).

### NMR Detection of Target Proteins in Lysates

Significant line-broadening persisted in the C2α spectrum even after the cell lysis (**Figure S1C**). However, addition of both EDTA and RNase A resulted in the appearance of well-dispersed ^15^N-^1^H_N_ cross-peaks indicative of a fully folded protein (**Figure 2D**). The line-broadening can therefore be attributed to the C2α interactions with metal ions and RNA^[20]^ in the crowded environment. The Ca^2+^-binding site of C2α resides on the apical CMBL region (**Figure 1C**) that can potentially interact with abundant divalent metal ions such as Mg^2+^. Indeed, only RNase A treatment was required to recover full spectra when C2α-expressing cells were grown in the presence of 50 μM EDTA and reduced Mg^2+^ levels. We hypothesized that the C2α region responsible for RNA interactions is the electropositive lysine-rich cluster, or LRC (**Figures 2A, S2A**) that mediates PKC recruitment to membranes through interactions with negatively charged lipids such as phosphatidylserine (PtdSer) and phosphatidylinositol-4,5-bisphosphate (PtdIns4,5P_2_).^[16a, 21]^ When we mutated two of the four Lys residues of the LRC to Glu, EDTA-only treatment was sufficient to obtain a well-resolved spectrum (**Figure S2A-B**) indicating that interactions with RNA are attenuated in this charge-reversal KEKE C2α mutant. In contrast to C2α, the lysate of cells expressing C1Bα did not require addition of RNase A or EDTA to produce a well-dispersed chemical shift pattern that reports on the fully folded protein (**Figure 2E**).

Pairwise comparison of the *in-cell* and lysate spectra for C1Bα and C2α (**Figures 2A-D, 2B-E, S1D**) suggests that line-broadening in the *in-cell* data is caused by both proteins engaging in significant quinary interactions with intracellular components. The propensity to engage in quinary interactions correlates with the existence of distinct electropositive regions, as both proteins have an isoelectric point of 8.7. In C1Bα, the relevant region is likely to be the α1 helix that is responsible for PtdSer recognition at the membranes (**Figure 2E**).^[15a]^ In C2α it is the CMBLs and the LRC regions whose interactions with intracellular binding partners might be enhanced in much the same way as Ca^2+^ binding facilitates the PKC interactions with negatively charged phospholipids. We note that in contrast to C1Bα and C2α, α_3_DIV is acidic (pI=5.8) and its electrostatic surface presents a rather disperse distribution of positively and negatively charged regions. As a result, α_3_DIV does not engage in any significant quinary interactions, evident by the comparable quality of the *in-cell* and lysate spectra (**Figure 2C-F**). Collectively, our data recommend the use of lysates to probe Cd^2+^ interactions with PKCα regulatory domains, and intact cells to probe Cd^2+^ interactions with α_3_DIV.

### Cd^2+^ Substitutes for Zn^2+^ in C1Bα Without Loss of Ligand-sensing Function

Structural Zn^2+^ ions, each coordinated in a tetrahedral geometry by a Cys_3_His motif are essential for the stability of the C1Bα fold (**Figure 3A**).^[15a]^ The affinity of Zn^2+^ to thiol-rich structural protein sites is estimated to be in the pico-to-femtomolar range. To determine if C1Bα incorporates Cd^2+^ in the presence of native Zn^2+^, we expressed the protein with 25 μM Zn^2+^ and 50 μM Cd^2+^ added to the *E. coli* culture. The [^15^N-^1^H] SOFAST-HMQC spectrum of cell lysate differs from that obtained in the Zn^2+^-only control experiment (**Figure 3A**). Notably, residues located at and near the Zn^2+^ coordination sites 1 and 2 show additional subset of amide cross-peaks whose position matches those previously obtained for the Cd^2+^-treated protein in an aqueous buffer.^[8e]^ We conclude that in the crowded environment Cd^2+^ successfully competes with Zn^2+^ for the Cys_3_His motifs of C1Bα and supports its three-dimensional fold.

**Figure 3.**
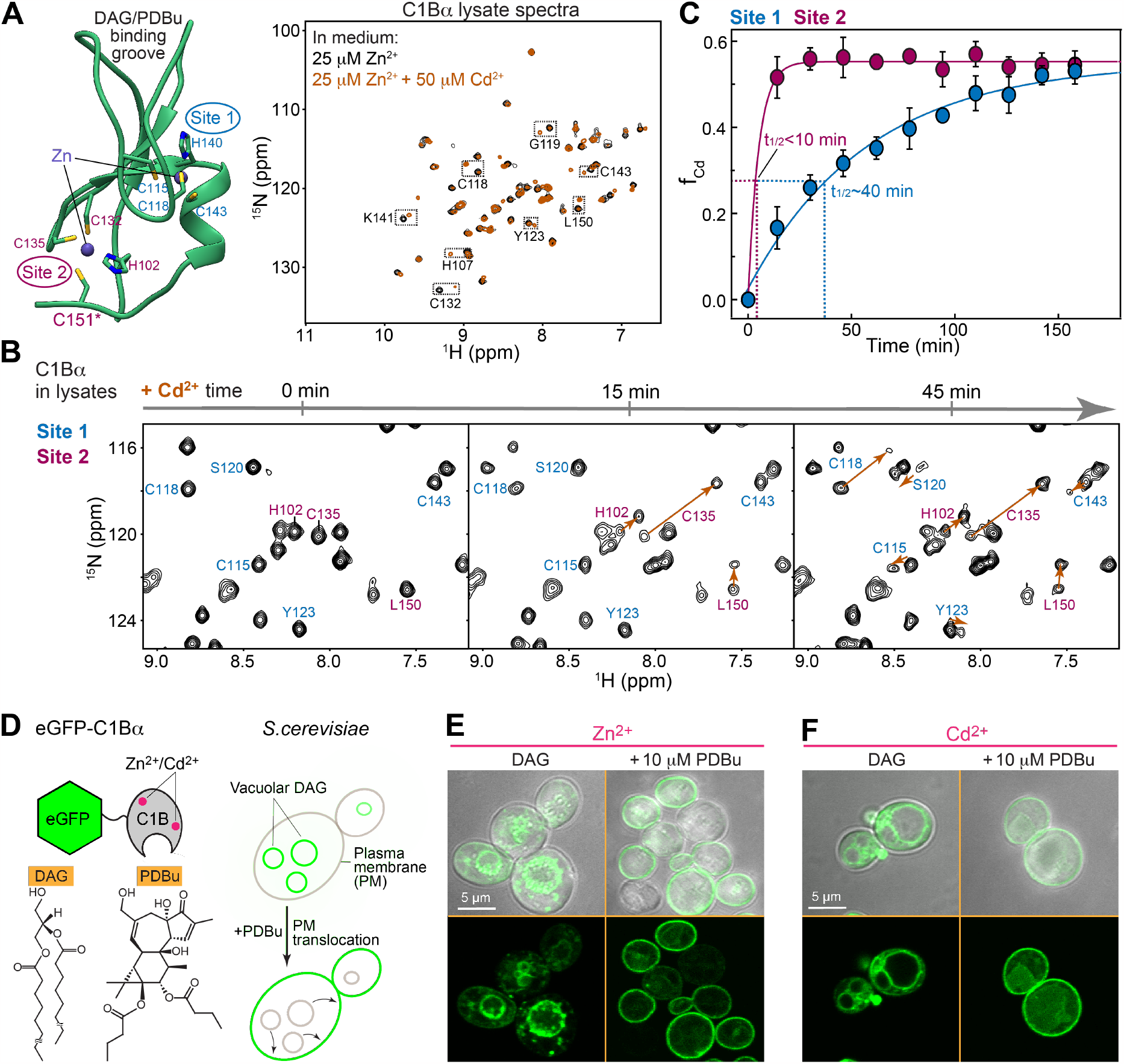
Displacement of structural Zn^2+^ by Cd^2+^ generates a functionally competent C1Bα fold. (A) Cys_3_His structural Zn^2+^ sites of C1Bα (left panel, PDB:2ELI) and the spectroscopic signatures of their Zn^2+^→Cd^2+^ substitution (right panel). The latter is illustrated by the superposition of [^15^N-^1^H] SOFAST-HMQC lysate spectra of cells expressing C1Bα in the presence of Zn^2+^ (black) and Zn^2+^-Cd^2+^ mixture (brown). Cd^2+^ binding to C1Bα results in the appearance of distinct amide resonances for several residues (highlighted by squares). (B,C) Kinetics of Zn^2+^→Cd^2+^ substitution initiated by the addition of 2-fold molar excess of Cd^2+^ to Zn^2+^-bound C1Bα. (B) shows expansions of representative [^15^N-^1^H] SOFAST-HMQC lysate spectra collected at 3 time points. The cross-peak labels of residues responsive to Cd^2+^ binding at sites 1 and 2 are color-coded blue and maroon, respectively. The arrows connect cross-peaks that belong to the same residue in Zn^2+^- and Cd^2+^-bound C1Bα species. (C) Time course of Zn^2+^→Cd^2+^ substitution reported by the mean fractional population of Cd^2+^ bound C1Bα (f_Cd_). Solid lines are to guide the eye. t_1/2_ are estimated as the time needed to reach 50% of the plateau f_Cd_ values. (D) Chemical structures of C1Bα ligands and experimental design of fluorescence microscopy experiments in S. cerevisiae. C1Bα is fused with eGFP to enable detection of intracellular DAG-dependent localization and translocation to plasma membrane upon stimulation with phorbol 12,13-dibutyrate (PDBu). (E,F) Live-cell confocal microscopy images of S. cerevisiae cells grown in SD medium (-URA) supplemented with either 10 μM Zn^2+^ (E) or Cd^2+^ (F). The membrane localization pattern of eGFP-C1Bα is similar when Cd^2+^ is added to the medium instead of Zn^2+^.

To establish the potency of Cd^2+^ in displacing Zn^2+^ from C1Bα, we monitored the metal ion substitution reaction by NMR spectroscopy in real time. The experiment involved adding purified 0.2 mM Zn^2+^-containing [U-^15^N] enriched C1Bα to the concentrated lysate of *E. coli* with no expression plasmid; initiating the Zn^2+^→Cd^2+^ substitution reaction by adding two-fold molar excess of Cd^2+^; and collecting [^15^N-^1^H] SOFAST-HMQC spectrum every ∼15 minutes. We find that the formation of C1Bα species with Cd^2+^ at site 2, C1Bα_Cd(2)_, occurs within the first 15 minutes and is manifested by the appearance of characteristic amide cross-peaks of His102, Cys135, and Leu150 (**Figure 3B**). A considerably slower formation of C1Bα species with Cd^2+^ at site 1, C1Bα_Cd(1)_, is reported by the appearance of characteristic amide cross-peaks of Cys118, Cys143, and Cys115.

To evaluate the relative susceptibility of Zn^2+^-coordinating Cys_3_His motifs to Cd^2+^ substitution, we calculated the fractional populations f_Cd_ of C1Bα_Cd(1)_ and C1Bα_Cd(2)_ species using NMR peak intensities and analyzed their accumulation kinetics. Site 2 is considerably more susceptible to Cd^2+^ (t_1/2_ < 10 min) than site 1 (t_1/2_ ∼ 40 min) (**Figure 3C**). This agrees well with our previous findings that site 2 of C1Bα is dynamic and prone to oxidative modifications.^[22]^ The dynamics is associated with the loss of the Zn-S(Cys151) coordination bond (**Figure 3A**) and the concomitant local conformational change of the protein. The latter increases solvent exposure of the Cys132-Cys135-Cys151-His102 motif, making it more susceptible to Cd^2+^ invasion. Of note, both kinetic curves shown in **Figure 3C** plateau at the f_Cd_ values of ∼55%, suggesting that in lysates the affinities of sites 1 and 2 to Cd^2+^ are comparable.

How does Cd^2+^ substitution affect the lipid-sensing function of C1Bα? We chose two ligands to probe the C1Bα function in live cells: (i) *sn*-1,2-diacylglycerol (DAG), a native lipid activator of PKC; and (ii) phorbol 12,13-dibutyrate (PDBu), a potent tumor-promoting activator of PKC (**Figure 3D**). Our recent structural work on the C1::ligand complexes^[15b]^ demonstrates how the integrity of the C1B domain and particularly of its apical loop region (**Figure 3A**) are essential for the recognition and capture of its membrane-embedded ligands. To evaluate the effect of Cd^2+^ on the C1Bα function, we monitored the ligand-dependent localization of the eGFP-C1Bα fusion protein in live *S. cerevisiae* cells using fluorescence microscopy (**Figure 3D**). *S. cerevisiae* cells were chosen because they provide a eukaryotic environment with high basal level of endogenous DAG.

We find that in the presence of 10 μM Zn^2+^ eGFP-C1Bα preferentially localizes to vacuolar rather than plasma membrane DAG pools (the plasma membrane is marked by the cell boundaries in bright field images shown in **Figure 3E-F**). The membrane localization pattern is similar when Cd^2+^ is added to the medium instead of Zn^2+^, indicating that Cd^2+^-substituted C1Bα retains DAG sensitivity. Treatment of cells with PDBu, a high-affinity competitor of DAG, results in essentially quantitative translocation of eGFP-C1Bα to the plasma membrane. That response was identical for cells exposed to Zn^2+^ or Cd^2+^. The appearance of cytoplasmic puncta were observed only when cells were treated with higher concentrations of Cd^2+^ (25 μM in medium, **Figure S3**). These puncta were frequently associated with the limiting membrane of the vacuole, and likely correspond to the fraction of eGFP-C1Bα subspecies that misfold and aggregate. The properly folded subspecies remain ligand-responsive and translocate to the plasma membrane upon PDBu treatment (**Figure S3**).

In summary, we established that Cd^2+^ targets high-affinity structural Zn^2+^ sites of C1Bα leading to the formation of Cd^2+^-substituted C1Bα that retains its ligand-sensing function. To our knowledge, the only other reported case of “isofunctional” Cd^2+^ substitution of Zn^2+^ in proteins is the Cadmium carbonic anhydrase of marine diatoms, which retains its catalytic activity as a Cd^2+^-bound complex.^[23]^ Our findings indicate that C1 domains of other PKC isoforms as well as those of other diacylglycerol effector proteins are potential targets of Cd^2+^. The co-occurrence of Zn and Cd-folds in C1Bα is analogous to how metallothionines incorporate Cd^2+^ in mixed Zn-Cd thiolate clusters.^[24]^ Given that many transcription factors bind Zn^2+^ via zinc finger motifs, and that some of these DNA-binding proteins execute Zn^2+^ sensing functions, there is building evidence to indicate that Cd^2+^ substitution in these cases contributes to Cd^2+^ toxicity.^[25]^ In this regard, as Zn^2+^ binding is a key feature of a large number of proteins in addition to transcription factors, the potential target space for Cd^2+^ substitution is large.^[26]^

### C2α Interacts with Cd^2+^ in *E. coli* Lysates

C2α binds its native metal ion, Ca^2+^, via an all-oxygen coordination environment provided by 5 conserved aspartate residues of the CMBL region (**Figure 4A**). It can accommodate up to 3 Ca^2+^ ions whose affinities fall into the μM-mM range. Our previous work has established, using biophysical measurements and structural data, that in aqueous buffer solutions Cd^2+^ binds to C2α with comparable stoichiometry and >30-fold higher affinity.^[8b]^ Since oxygen is a harder Lewis base than sulfur, Cd^2+^ affinity to all-oxygen coordination sites is generally weaker than that to all-sulfur sites. To determine if C2α-Cd^2+^ interactions take place in the crowded molecular environment that contains many potential Cd^2+^ chelators, we conducted experiments on lysates of cells that express isotopically enriched C2α. To generate a clean metal-free reference state of C2α, we expressed the protein in the presence of 50 μM EDTA in the medium under limiting (10 μM) Mg^2+^ supplementation. The [^15^N-^1^H] SOFAST-HMQC spectrum of RNase A-treated lysate remained unchanged upon EDTA addition and showed well-dispersed cross-peaks characteristic of a fully folded protein (**Figure 4B**).

**Figure 4.**
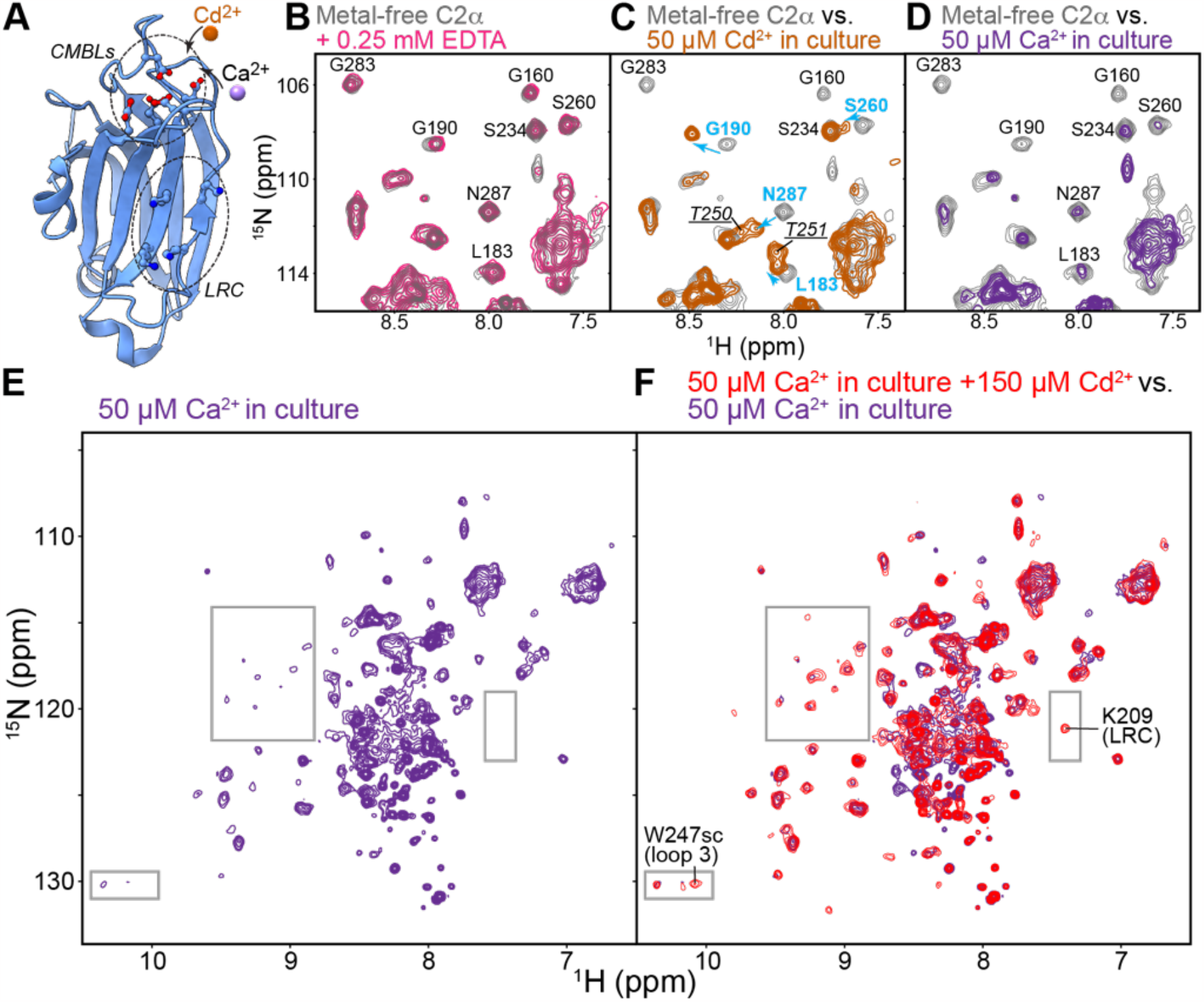
C2α binds Cd^2+^ and Ca^2+^ in cell lysates. (A) Three-dimensional structure of C2α (PDB:3GPE) showing the location of Ca^2+^- and membrane-binding loops (CMBLs) and the Lysine-Rich Cluster (LRC). CMBLs bind Cd^2+^, while LRC interacts with anionic species. (B-D) Expansions of [^15^N-^1^H] SOFAST-HMQC spectra collected on lysates of E. coli cells expressing C2α in the presence of 10 μM Mg^2+^ (B), 50 μM Cd^2+^ (C), and 50 μM Ca^2+^ (D) in the medium. All lysates were treated with RNase A prior to data collection. In (B), the C2α spectrum in Mg^2+^-containing lysate (gray) does not appreciably change upon EDTA treatment (pink), indicating that it corresponds to the metal-free state of the protein. In (C), several C2α residues in Cd^2+^-containing lysate (brown) show chemical shift changes characteristic of the Cd^2+^-bound protein. The changes are shown with blue arrows. The cross-peaks of the underlined residues T250 and T251 are broadened in the apo C2α but reappear upon Cd^2+^ binding. In (D), the C2α spectrum in Ca^2+^-containing lysate (purple) shows significant linebroadening compared to the metal-free protein (gray). (E) Full [^15^N-^1^H] SOFAST-HMQC spectrum of C2α in Ca^2+^-containing lysate (purple), and (F) its overlay with spectrum obtained upon addition of 150 μM Cd^2+^ into the lysate (red). Cd^2+^ addition results in the recovery of resonance intensities, evident in the spectral regions highlighted by rectangles. W247 and K209 are noted as representative residues for loop 3 and LRC regions, respectively.

Next, we prepared lysate samples of cells grown with either 50 μM Cd^2+^ or Ca^2+^ but no EDTA added in culture. The chemical shifts of the C2α amides in Cd^2+^-containing lysate clearly report on the formation of the C2α-Cd^2+^ complex^[8b]^ (**Figure 4C**). For instance, the chemical shift perturbations of Gly190, Asn287, and Leu183 amide resonances are similar to those detected in C2α upon Cd^2+^ binding in aqueous buffer.^[8b]^ The cross-peaks of Thr150 and Thr151 of CMBL3 that are exchange-broadened beyond detection in the metal-free state, are readily detectable in the Cd^2+^-containing lysate due to the rigidification of the CMBL region that takes place upon metal ion complexation (**Figure 4C**).

In sharp contrast with Cd^2+^, the C2α amide peaks in the Ca^2+^-containing lysate are significantly broadened (**Figures 4D-E**). Since this broadening affected most protein residues and not just those of the Ca^2+^-binding region, we conclude that Ca^2+^-complexed C2α engages extensively in quinary interactions. Indeed, Ca^2+^-induced NMR line broadening is reversible by EDTA treatment. When Cd^2+^ is added to the Ca^2+^-containing lysate many of the protein cross-peaks recover their intensities (**Figure 4F**). The latter finding suggests that Cd^2+^ can displace native Ca^2+^ from C2α and thereby affect its propensity to engage in quinary interactions. One plausible explanation for this are coordination preferences of Ca^2+^ and Cd^2+^. In biological macromolecules, Ca^2+^ has a potential to expand its coordination number up to 8, whereas for Cd^2+^ the coordination numbers of 4 to 6 are more common.^[27]^ Therefore, Ca^2+^ might directly mediate the interactions of C2α with polyanionic compounds within cells/lysates. This interpretation is supported by our previous findings that Ca^2+^-bound C2α and both C2 domains of Synaptotagmin 1 readily associate with anionic membranes (specifically anionic lipids PtdSer and PtdIns(4,5)P_2_) as part of their signal transduction mechanism, while Cd^2+^-bound proteins are unable to bind PtdSer-containing membranes.^[8b, 8c]^

In summary, we have demonstrated that Cd^2+^ is capable of targeting oxygen-rich sites and displacing native metal ion Ca^2+^ from C2α. In a broader context, aspartate and glutamate side chains frequently serve as bidentate ligands in the selectivity filters of ion channels and metal-ion sensing EF-hand proteins. The NMR-based approach presented here can be applied to test the Cd^2+^ susceptibility of such protein targets in the crowded environment.

### α_3_DIV Sequesters Cd^2+^ in *E. coli* Cells

α_3_DIV is a designed metal-ion sensing protein of 73 amino acids that contains a tri-cysteine motif (**Figure 5A**). This motif binds heavy metal ions Cd^2+^, Pb^2+^, and Hg^2+^ with sub-μM affinities in aqueous solutions.^[17]^ The metal-ion binding site of α_3_DIV resembles thiol-rich coordination sites found in bacterial MerR family of transcriptional regulators^[8a, 28]^ that play key roles in heavy-metal efflux and detoxification processes. While no high-resolution metal-ion bound structure of α_3_DIV is available, ^113^Cd NMR and ^111m^Cd PAC spectroscopy measurements suggest that Cd^2+^ adopts a pseudo-tetrahedral geometry CdS_3_(O/N)^-^, where O belongs to the exogenous water molecule and N belongs to His72 of α_3_DIV.^[17]^ We sought to evaluate the potential of α_3_DIV as a Cd^2+^-sensing and sequestering protein in the cellular environment.

**Figure 5.**
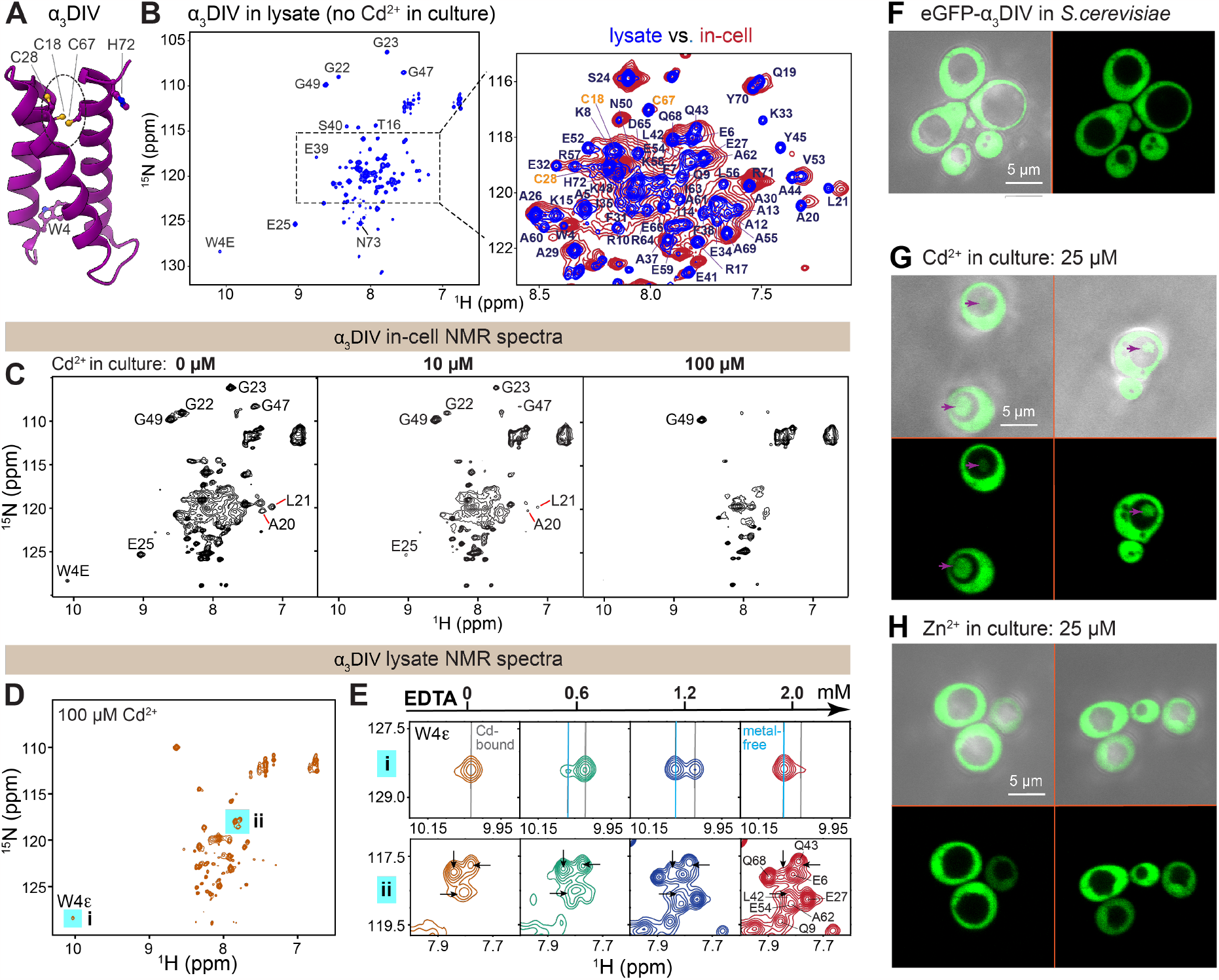
α_3_DIV sequesters Cd^2+^ in cells. (A) Three-dimensional structure of α_3_DIV (PDB:2MTQ) highlighting Cd^2+^ coordination motif formed by three Cys residues: 18, 28, and 67. His72 may also be involved in metal ion coordination. (B) [^15^N-^1^H] SOFAST-HMQC spectrum of α_3_DIV in E. coli Cd^2+^-free lysate (blue, left panel) and its overlay with the in-cell spectrum (red, right panel). (C) In-cell [^15^N-^1^H] SOFAST-HMQC spectra of α_3_DIV at 0, 10, and 100 μM Cd^2+^ added in culture. Progressive line-broadening is observed upon increasing Cd^2+^ concentration. (D) [^15^N-^1^H] SOFAST-HMQC spectrum of α_3_DIV in lysate generated from cells grown with 100 μM Cd^2+^ in culture. Addition of increasing EDTA concentration to this sample results in the recovery of apo α_3_DIV resonances, as shown by the adjacent spectral expansions of regions i and ii in (E). (F) Live-cell confocal microscopy image of S. cerevisiae cells expressing eGFP-α_3_DIV grown in SD medium (-URA). The protein shows a diffuse cytosolic localization pattern. (G) Addition of Cd^2+^ to the growth medium results in the import of detectable eGFP-α_3_DIV fraction (marked with pink arrows) into the vacuolar lumen. (H) Addition of Zn^2+^ has no significant effect on the cytosolic distribution of eGFP-α_3_DIV. In (F-H), bright field images mark the cell boundary.

The ability of α_3_DIV to bind heavy metal ions relies on the pre-organization of the tri-cysteine motif through the formation of the triple-helix bundle. This assembly process is pH dependent.^[18]^ To evaluate the α_3_DIV state in the crowded environment, we mapped the amide cross-peak assignments obtained in aqueous buffer^[18]^ onto the [^15^N-^1^H] SOFAST-HMQC spectra of Cd^2+^-free lysates and intact cells expressing α_3_DIV (**Figures 5B, C**). Based on the excellent agreement between the α_3_DIV chemical shifts in all three environments (buffer, lysate, and intact cells), we conclude that α_3_DIV maintains its folded state with the pre-organized tri-cysteine motif in the cellular milieu.

To determine if α_3_DIV can serve as a cellular reporter of Cd^2+^ uptake, we expressed the protein in *E. coli* with 10 and 100 μM Cd^2+^ added in culture. Comparison of the *in-cell* NMR spectra of α_3_DIV at 0, 10 and 100 μM Cd^2+^ shows significant peak broadening as Cd^2+^ concentration increases (**Figure 5C**). Although we cannot exclude the engagement of α_3_DIV in quinary interactions, the most likely cause is the chemical exchange of Cd^2+^-bound α_3_DIV that is intermediate on the chemical shift timescale. In fact, a similar pattern of line-broadening was observed previously in the NMR spectra of Pb^2+^- and Hg^2+^-bound α_3_DIV (aqueous buffer), and was attributed to structural dynamics associated with metal ions adopting distinct coordination geometries within the asymmetric tri-cysteine site.^[18]^ Not surprisingly, Cd^2+^-induced peak broadening persists in cell lysates (**Figures 5D and S4**). Treatment of lysates with increasing concentrations of EDTA removes bound Cd^2+^ from α_3_DIV and results in the recovery of resonances that correspond to the metal-free α_3_DIV (**Figures 5E, S4A-B**). This is aptly illustrated by the behavior of the ^15^Ne-^1^He Trp4 cross-peak that has distinct ^1^H chemical shifts for the apo and Cd^2+^-bound α_3_DIV species (**Figures 5E, S4C**). Together, these findings demonstrate that α_3_DIV sequesters Cd^2+^ taken up by *E. coli*.

Next, we tested the specificity of α_3_DIV as a Cd^2+^ sensor in both prokaryotic and eukaryotic environments by introducing Zn^2+^ as a competing metal ion. The NMR spectrum of cells grown with 100 μM Zn^2+^ added in culture revealed only insignificant changes compared to the no-metal ion control, suggesting that no specific α_3_DIV-Zn^2+^ interactions take place in the cellular environment (**Figure S5A**). Very high concentrations of Zn^2+^, 0.4 mM, added externally to cell lysates were required to detect minor changes in the NMR spectra (**Figure S5B**), further supporting our conclusion that no specific high-affinity interactions with Zn^2+^ take place.

We assessed the Cd^2+^- and Zn^2+^-dependent behavior of α_3_DIV in eukaryotic environment by monitoring the localization of the eGFP-α_3_DIV in *S*.*cerevisiae* using fluorescence microscopy. In the absence of direct metal ion supplementation, eGFP-α_3_DIV shows a cytosolic distribution (**Figure 5F**). Upon addition of Cd^2+^ to the concentration of 25 μM, however, a detectable fraction of eGFP-α_3_DIV is imported into to the vacuolar lumen (**Figure 5G**). This relocalization is not observed upon addition of 25 μM Zn^2+^ (**Figure 5H**). We interpret these data to indicate that Cd^2+^-bound α_3_DIV is a substrate for direct transport from the cytoplasm into the vacuole via the cytoplasm-to-vacuole targeting (Cvt) pathway.^[29]^ We speculate eGFP-α_3_DIV has a propensity to aggregate and that it is this aggregation that induces its clearance from the cytoplasm.

In summary, we have demonstrated that α_3_DIV responds to cellular Cd^2+^ levels by showing characteristic NMR signatures and localization patterns. A combination of robust structural framework provided by the α_3_D triple-helix bundle and the engineered metal-ion binding sites provides an opportunity to generate a variety of metal sensors and chelators.^[30]^ Potential applications of these protein scaffolds include bioremediation using microbial systems, investigation of heavy metal resistance mechanisms, and rational design of coordination environments to achieve high metal selectivity.^[31]^ *In-cell* NMR and direct imaging methods, as demonstrated for the α_3_DIV-Cd^2+^ system, show significant promise in the evaluation of metal-ion binding properties in the crowded environments and in guiding further design efforts.

## Conclusions

Metal binding properties of proteins and peptides form the chemical foundation of many regulatory processes in the cell. These processes are of direct relevance to the application of metal ions for medicinal^[32]^ and biorthogonal^[33]^ purposes. Xenobiotic metals such as Cd^2+^ target proteins that rely upon nutritive divalent cations for structural and functional integrity. This feature defines the basis for why xenobiotic metals are so deleterious to living cells. From this perspective, characterization of protein interactions with toxic metals in cellular contexts is required for reliable identification of physiologically relevant targets, development of strategies for bioremediation, and creation or refinement of protein-based sensors.

In this work, we applied NMR spectroscopy to probe site-specific response of three proteins to Cd^2+^ binding in crowded environments provided by intact cells and their lysates. *E. coli* was the experimental model of choice for these studies given its high tolerance to toxic metal ions and the efficiency with which recombinant proteins can be labeled with stable isotopes. Cd^2+^ and the nutritive metals Ca^2+^ and Zn^2+^ were introduced during the window of target protein synthesis and thereby provided a reliable means for generating distinct metalated target protein states in cellular contexts. A major conclusion of our work is that Cd^2+^ targets sulfur-rich, oxygen-rich, as well as mixed sulfur/nitrogen protein coordination sites. That this heavy metal ion successfully competes with native metal ions Ca^2+^ and Zn^2+^ is a surprising result given Cd^2+^ availability is expected to be limited in the highly chelating environment of the cytoplasm. We also demonstrate that Cd^2+^ binding event is associated with distinct protein chemical shift perturbations and modulation of quinary interactions. The distinct spectroscopic signatures associated with metal-ion binding events attest to the potential of NMR spectroscopy as a general tool to probe the interactions of proteins and peptides with toxic metals in crowded biomolecular environments. Moreover, we show that augmentation of NMR experiments with live cell imaging represents a particularly powerful approach that can be extended to other xenobiotic metal ions.

## Acknowledgements

This work was supported by the National Science Foundation (NSF) grant CHE-1905116 and National Institute of Health (NIH) grant R01 GM108998 to T.I.I. We acknowledge Dr. Robert Burghardt for assistance with data collection on the Zeiss LSM 780 NLO multiphoton microscope equipped with Airyscan detector (Texas A&M Image Analysis Laboratory, School of Veterinary Medicine & Biomedical Sciences). We thank Ms. Savana M. Green for assistance with handling yeast cultures and collection of fluorescence imaging data, and Mr. Nowlan Savage for his contribution to the development stages of the project. We acknowledge Dr. Vytas A. Bankaitis (Texas A&M University) for guidance on the interpretation of yeast imaging data and for critical reading of the manuscript.

## Author contributions

S.S.K. and T.I.I conceptualized the project and designed the experiments. S.S.K. prepared the samples, collected and analyzed the NMR data, and confocal fluorescence imaging datasets. S.S.K. and T.I.I. wrote the manuscript. T.I.I. supervised the project.

