## Supporting Information for "Protein-Cadmium Interactions in Crowded Biomolecular Environments Probed by In-cell and Lysate NMR Spectroscopy"

**Table of Contents**

1. Experimental Procedures……………………………………………………..3-6

1.1 Materials

1.1.1 *E. coli* expression constructs

1.1.2 *S. cerevisiae* expression constructs

1.1.3 Metal stock solutions, buffer components, ligands, and isotopes

1.2 Methods

1.2.1 NMR Spectroscopy

1.2.2 NMR sample preparation procedures

1.2.3 C1Bα-specific procedures

1.2.4 C2α-specific procedures

1.2.5 $\alpha_3$DIV-specific procedures

1.2.6 Fluorescence microscopy in *S. cerevisiae*

2. Results and Discussion…………………………………………………………7-11

Figure S1
Figure S2
Figure S3
Figure S4
Figure S5

3. Supplementary References……………………………………………………….12

### Experimental Procedures

#### 1.1 Materials

##### 1.1.1 *Escherichia coli* expression constructs

The cDNA fragments (Open Biosystems, GE) encoding the PKCα C1B domain (C1Bα, residues 100-152, *Mus musculus*) and C2 domain (C2α, residues 155-293, *Rattus norvegicus*) were cloned into a pET-SUMO expression vector (Invitrogen). Cloning, overexpression in *E. coli* BL21(DE3) cells, and purification procedures are described elsewhere.[1] To express these proteins for *in-cell* and lysate NMR experiments, the 6xHis-SUMO solubility tag was deleted using Q5™ site-directed mutagenesis kit (NEB Inc.). The C2α "KEKE" variant was prepared by mutating K197 and K209 residues to glutamic acid using the same kit. The $\alpha_3$DIV synthetic gene subcloned into the pET-15b expression vector was obtained from GenScript®.

##### 1.1.2 *Saccharomyces cerevisiae* expression constructs

Wild-type *S. cerevisiae* strain CTY182 (*MATα ura3-52 lys2-801 his3Δ-200*) was used in all experiments. The DNA fragments encoding C1Bα and $\alpha_3$DIV were PCR-amplified from the corresponding bacterial expression vectors with the addition of NotI (at 5' end) and HindIII (3' end) restriction sites. The resulting PCR products were digested with NotI and HindIII, and cloned into the YC-type (centromeric) expression vector with eGFP as the N-terminal fusion tag. The final constructs express the eGFP-C1Bα and eGFP-$\alpha_3$DIV under constitutive tetO7-CYC1 promoter.

##### 1.1.3 Metal stock solutions, buffer components, ligands, and isotopes

The 100 mM stock solution of $Cd^{2+}$ was prepared by dissolving $Cd(NO_3)_2\cdot 4H_2O$ (Sigma-Aldrich) in HPLC-grade water. The working solutions of $Zn^{2+}$ and $Ca^{2+}$ were prepared from 100 mM stock solution of $ZnSO_4$ (Fluka® Analytical) and 1 M stock solution of $CaCl_2$ (Fluka® Analytical), respectively, in HPLC water. Unless specified otherwise, all samples for *in-cell* and lysate NMR experiments were made in 10 mM HEPES "NMR buffer" at pH 7.2, 75 mM KCl, 1 mM tris (2-carboxyethyl) phosphine (TCEP), and 8% $D_2O$. To remove the trace divalent cations, the buffer was treated with Chelex® 100 matrix (sodium form, dry; Sigma-Aldrich) using the batch method. The C1Bα ligand, phorbol 12,13-dibutyrate (PDBu), was purchased from Sigma-Aldrich and dissolved in DMSO for addition to *S. cerevisiae* cultures. Isotopic media components: $^{15}NH_4Cl$ (99%), $D_2O$ (99.8 %), and the rich bacterial growth supplement Bioexpress® 1000 [U-D 98%, U-$^{15}$N 98%] were obtained from Cambridge Isotope Laboratories, Inc.

#### 1.2 Methods

##### 1.2.1 NMR spectroscopy

The NMR experiments were conducted on the Bruker Avance III HD spectrometers operating at the $^1$H Larmor frequencies of 800 MHz (18.8 T) and 600 MHz (14.1 T), both equipped with cryogenically cooled probes; and the 500 MHz (11.7 T) spectrometer equipped with a room temperature probe. The sample temperature was calibrated to 298.15 $^{0}$K using D4-98% Methanol. 2D heteronuclear chemical shift correlation spectra, ~15 minutes each, were collected with the $^{15}N$-$^{1}H$ SOFAST-HMQC scheme[2], processed with nmrPipe[3], and analyzed with NMRFAM-Sparky.[4]

#### 1.2.2 NMR sample preparation procedures

The workflow for the preparation of *in-cell* and lysate NMR samples of the proteins is shown in Figure 1A. Briefly, *E. coli* BL21 (DE3) cells were transformed with the plasmids encoding the protein of interest and cultured in 200 ml of LB broth supplemented with 50 mg/ml kanamycin (for C1B$\alpha$ and C2$\alpha$) or 100 mg/ml ampicillin (for $\alpha_3$DIV). Upon reaching $OD_{600}$ of ~0.4-0.6, the cells were harvested by centrifugation at 4,000 rpm for 20 min, washed with 20 ml of M9 salts, and re-suspended in 100 ml of M9 minimal medium (prepared in $D_2O$) containing 1 ml Bioexpress® 1000 [U-D 98%, U-$^{15}N$ 98%], 0.1 g $^{15}NH_4Cl$ (99%), 1 ml of Kao-Michayluk vitamin solution (100x; Sigma). The growth conditions, metal ion supplementation, and induction of expression were protein-specific and are detailed in the sections below. To prepare samples for *in-cell* NMR spectroscopy, 100 mL of cell culture was centrifuged at 1000 x g for 10 min. The pellet was washed once with 1.2 mL of the “NMR buffer”, and resuspended in the same buffer to form a 50% v/v slurry.

After the completion of *in-cell* NMR measurements, the samples were subjected to gentle centrifugation at 1000 x g for 5 min. The supernatant was used as the protein leakage control. The cell pellet was used to prepare lysate samples. Those were generated by (i) resuspending cell pellets in 1.2 ml of “NMR buffer” supplemented with 2.5 μl Lysozyme (5 mg/ml), a flake of DNAse, and 2.5 μl HALT™ protease inhibitor cocktail (100x); (ii) sonication; and (iii) clarification of the resulting lysate solution by centrifugation at 13,000 rpm for 10 min.

#### 1.2.3 C1Bα-specific procedures

To prepare $Zn^{2+}$- and $Cd^{2+}$-containing NMR samples of [$^{2}H$,$^{15}N$]-C1B$\alpha$, the cell pellets from the LB cultures were re-suspended in M9 minimal medium (composition described in 1.2.2) that was additionally supplemented with either 25 μM $Zn^{2+}$ or a mixture of 25 μM $Zn^{2+}$/50 μM $Cd^{2+}$. The cell cultures were equilibrated for 1 hour at 16 $^{0}$C, and the protein expression was induced with 0.5 mM IPTG for 12 hrs. Preparation of samples for *in-cell* NMR spectroscopy was as described in 1.2.2. To make lysate samples, the sonicated cell pellet suspension was diluted two-fold, subjected to ultracentrifugation at 50,000 rpm, and brought down to the 0.6 mL volume by concentrating with a 3 kDa cutoff filter.

For the measurements of $Zn^{2+}$ displacement kinetics, purified 0.2 mM [U-$^{15}N$] C1B$\alpha$ was added to the lysate generated from untransformed *E. coli* BL21(DE3) cells and the reference $^{15}N$-$^{1}H$ SOFAST-HMQC spectrum was recorded. The displacement reaction was initiated by adding 0.4 mM $Cd^{2+}$ to the sample and monitored in real time by collecting ~15 min $^{15}N$-$^{1}H$ SOFAST-HMQC spectra. The residue-specific fractional population of Cd-bound species, $f_{Cd}$, was calculated as:

$$f_{Cd} = \frac{I_{Cd}}{I_{Cd} + I_{Zn}}$$

where $I_{Cd}$ and $I_{Zn}$ are the intensities of amide cross-peaks corresponding to the $Cd^{2+}$- and $Zn^{2+}$-bound C1Bα species, respectively. The mean values of $f_{Cd}$ were calculated using six residues as reporters for Site 1: C115, H117, S120, Y123, K141, C143; and five residues as reporters for Site 2: K103, C135, L150, C151, G152. The error bars in Figure 3C are the standard deviations from the mean.

#### 1.2.4 C2α-specific procedures

To prepare the *in-cell* NMR samples of [$^{2}H$,$^{15}N$]-C2α or its KEKE mutant, the cell pellets were re-suspended in M9 minimal medium (composition described in 1.2.2) and equilibrated at 37 $^{0}$C for 20 min. For protein expression under limited $Mg^{2+}$ supplementation, the concentration of $MgSO_4$ in the M9 medium was reduced from 2 mM to 10 μM, and 50 μM EDTA was added after induction of protein expression with 0.5 mM IPTG for 3 hrs. For the $Cd^{2+}$/$Ca^{2+}$-containing samples, the respective metal ions were added to the M9 medium at 50 μM final concentration after induction, while keeping $Mg^{2+}$ at 10 μM. Preparation of samples for *in-cell* and lysate NMR spectroscopy was as described in 1.2.2. 0.25 mM EDTA and 5 μl of RNAse A (10 mg/ml, EMD Millipore) were added where appropriate. To detect $Ca^{2+}$ displacement by $Cd^{2+}$, $^{15}N$-$^{1}H$ SOFAST-HMQC spectra were collected before and one after addition of 0.15 mM $Cd^{2+}$ to the lysate of *E. coli* expressing [$^{2}H$,$^{15}N$]-C2α and grown with 50 μM $Ca^{2+}$ in culture.

#### 1.2.5 Preparation of $\alpha_3$DIV-containing NMR samples

To prepare the *in-cell* NMR samples of [$^{2}H$,$^{15}N$]-$\alpha_3$DIV, the cell pellets were re-suspended in M9 minimal medium (composition described in 1.2.2) and equilibrated at 37 $^{0}$C for 20 min. $Cd^{2+}$/$Zn^{2+}$-bound states of $\alpha_3$DIV were generated by supplementing the M9 medium with either 10 or 100 mM of $Cd^{2+}$/$Zn^{2+}$ prior to induction. Protein expression was induced with 0.5 mM IPTG for 2 hrs. Preparation of samples for *in-cell* and lysate NMR spectroscopy was as described in 1.2.2. To verify reversible, site-specific $Cd^{2+}$ chelation by $\alpha_3$DIV, the lysates generated from cells grown in the presence of 10 or 100 μM $Cd^{2+}$ were subjected to progressive additions of EDTA. At each EDTA concentration, an $^{15}N$-$^{1}H$ SOFAST-HMQC spectrum was collected to monitor the sequestration of protein-bound $Cd^{2+}$ ion by EDTA. To probe $Zn^{2+}$ interactions with $\alpha_3$DIV, 0.4 mM $Zn^{2+}$ was added to the lysates prepared from cells grown in the absence of $Zn^{2+}$. An $^{15}N$-$^{1}H$ SOFAST-HMQC spectrum was collected on the sample to assess the spectroscopic signatures of $Zn^{2+}$ association, followed by the addition of 0.4 mM EDTA to verify the reversible nature of the interaction.

#### 1.2.6 Fluorescence microscopy in *S. cerevisiae*

All *S. cerevisiae* cell cultures expressing eGFP-C1Bα and eGFP-α3DIV were grown overnight in synthetic-defined (SD) minimal medium (-URA) at 30 $^{0}$C with shaking at 250 rpm. The pH of the SD medium was set to 6.6 (as opposed to the usual pH of ~5) to facilitate the folding of $\alpha_3$DIV. Cells were prepared for confocal imaging by growing them either in the absence of explicit metal ion additive, or in the presence of either 25 μM $Zn^{2+}$ or $Cd^{2+}$. In addition, cells expressing eGFP-C1Bα were grown at either 10 μM $Zn^{2+}$ or $Cd^{2+}$. A notable growth lag of ~10 hrs (to reach the exponential phase) was observed for the $Cd^{2+}$-containing cultures. The cells were harvested at the exponential phase for imaging.

The confocal images were collected on the Zeiss LSM 780 NLO multiphoton microscope equipped with Airyscan detector using 63x oil immersion objective lens with numerical aperture of 1.44. For the acquisition of fluorescence signal, samples were excited at 488 nm with a detector range set to 490-560 nm (eGFP). The bright field images were collected using the transmitted light PMT detector. Live yeast cells were immobilized on the slides with Concanavalin A (GE Healthcare 17-0450-01) for all acquisitions. Post-acquisition processing of the images (collected as Z-stacks) was done using ImageJ2 (Fiji v 2.9.0/1.53t).[5] The figure panels were assembled in Adobe Illustrator.

### Results and Discussion

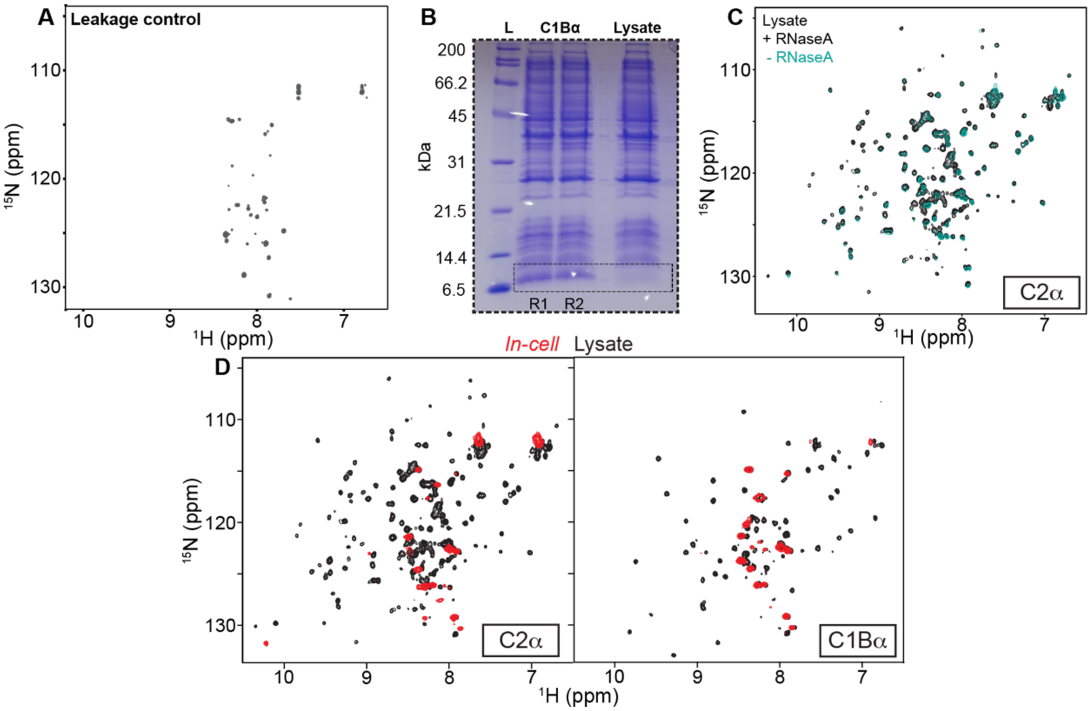


***Figure S1.*** *(A) Representative $^{15}$N-$^{1}$H SOFAST-HMQC spectrum collected as a protein leakage control. The spectrum shows resonances corresponding to the secreted bacterial metabolites. (B) Composition of the lysate samples of E. coli expressing C1Bα. The two independent replicates R1 and R2 (labeled "C1Bα") were diluted 4-fold for the SDS-PAGE detection. Lysate of cells without C1Bα (labeled "Lysate") is shown for comparison. (C) $^{15}$N-$^{1}$H SOFAST-HMQC spectral overlay of C2α in cell lysates treated with EDTA, with (black) and without (green) RNase A treatment. (D) $^{15}$N-$^{1}$H SOFAST-HMQC spectral overlays of C2α and C1Bα comparing their respective in-cell spectra with the lysate counterparts (both shown separately in Figure 2C-D for C2α and Figure 2E-F for C1Bα).*

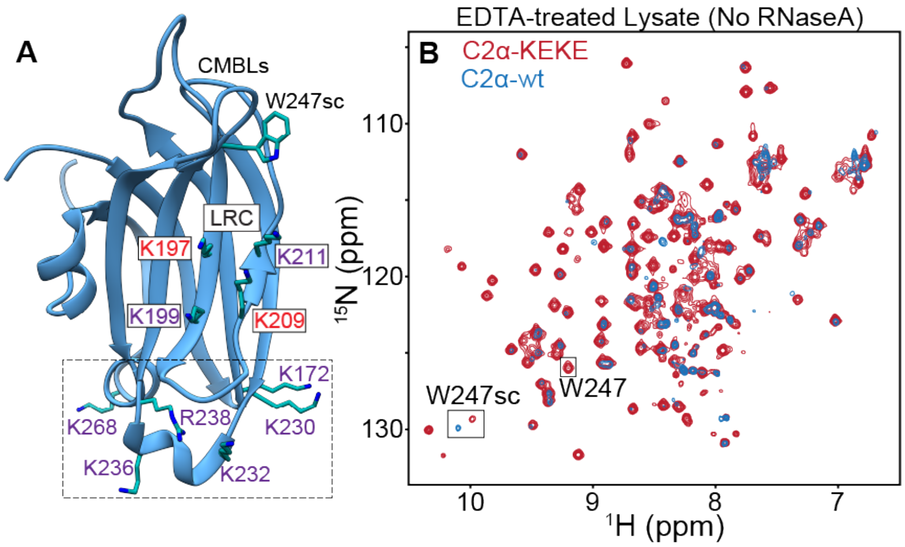


***Figure S2.*** *C2α-KEKE variant shows improved resonance dispersion and lysate detectability compared to the wild-type protein. (A) 3D representation of C2α (PDB ID: 3GPE, metal ions are not shown for clarity) showing $Ca^{2+}$ binding loops (CMBLs), the lysine-rich cluster (LRC), and the bottom region with additional basic residues. Among the four Lys residues of the LRC, K197 and K209 were mutated to glutamate to generate the "KEKE" variant. (B) [$^{15}N$-$^{1}H$] SOFAST-HMQC spectral overlays of C2α-KEKE (red) and wtC2α (blue) in lysates. The lysates were treated only with 0.25 mM EDTA, but not with RNase A. The side chain resonance of one of the CMBL-LRC junctional residue, W247, is perturbed in addition to the global intensity differences.*

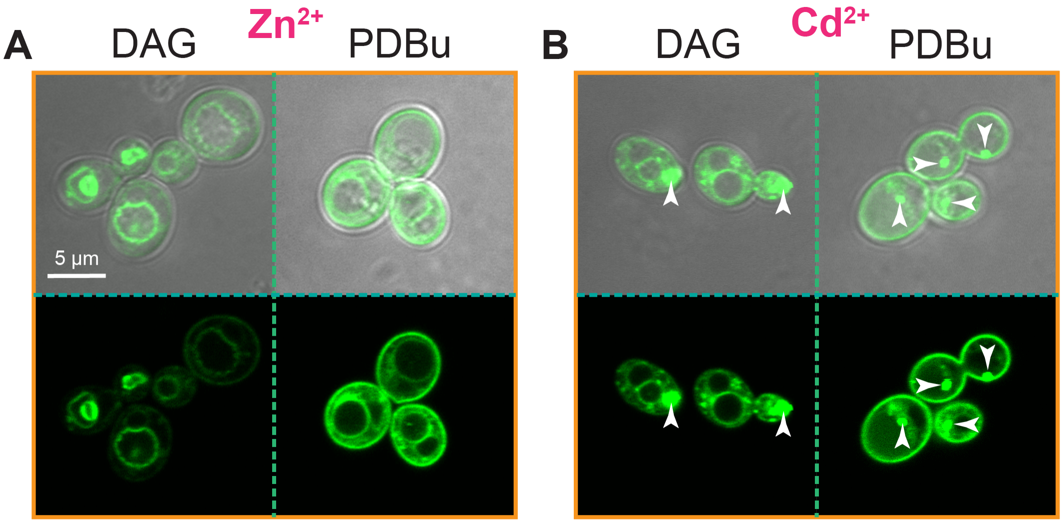


***Figure S3.*** *High concentrations of $Cd^{2+}$ result in the formation of prominent eGFP-C1B$\alpha$ puncta. Live-cell confocal microscopy images of eGFP-C1B$\alpha$ in yeast cells grown in SD medium (-URA) supplemented with either 25 $\mu$M $Zn^{2+}$ (A) or $Cd^{2+}$ (B). The top rows in (A) and (B) correspond to the bright field images. In (B), examples of large puncta in both the DAG and PDBu panels are marked by arrows. These substructures likely correspond to misfolded protein aggregates.*

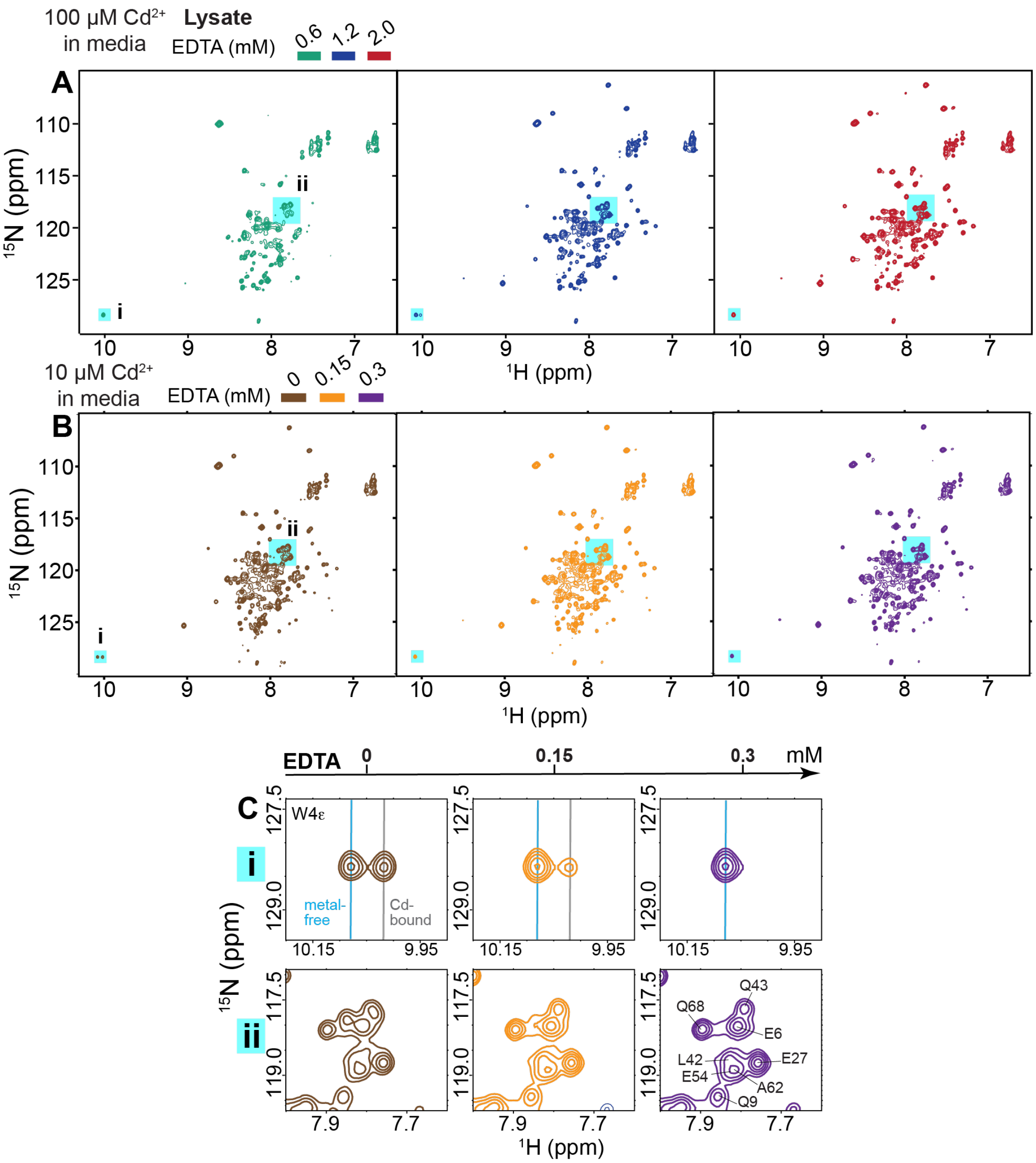


***Figure S4.*** *$Cd^{2+}$ sequestration by α₃DIV is reversible and concentration-dependent. (A) and (B) $^{15}N$-$^{1}H$ SOFAST-HMQC spectra of α₃DIV in E. coli lysate at different concentrations of EDTA. The samples were generated from cells exposed to 100 (A) and 10 (B) μM $Cd^{2+}$ in the growth medium. EDTA was added after the lysate preparation. (C) Spectral expansions of the boxed regions in (B). Addition of EDTA to the sample generated from cells exposed to 10 μM $Cd^{2+}$ results in the recovery of resonance intensities and full recovery of $Cd^{2+}$-free α₃DIV state, as reported by the chemical shifts. The equivalent spectral expansions for the 100 μM $Cd^{2+}$ sample are shown in Figure 5E.*

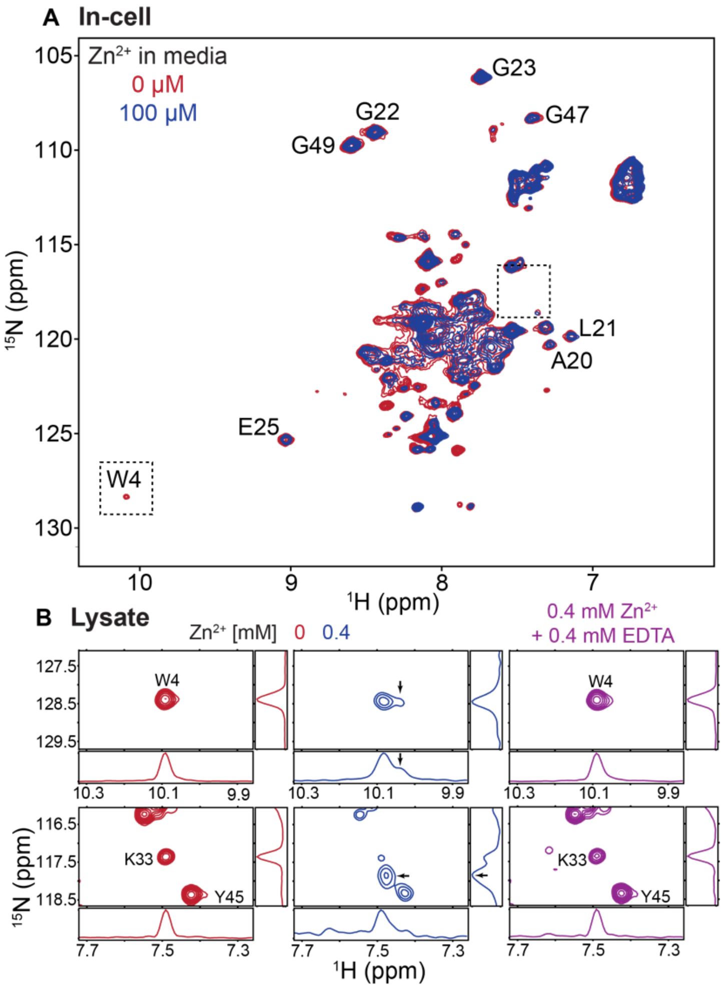


***Figure S5.*** *α₃DIV engages in weak non-specific interactions with $Zn^{2+}$. (A) In-cell $^{15}N$-$^{1}H$ SOFAST-HMQC spectra of α₃DIV in absence (red) and presence (blue) of 100 μM $Zn^{2+}$ in the growth medium. (B) Spectral expansions corresponding to the boxed regions of (A) in the absence of $Zn^{2+}$ (red), in the presence of 0.4 mM $Zn^{2+}$ (blue), and in the presence of 0.4 mM $Zn^{2+}$/0.4 mM EDTA (purple). The pattern of resonance perturbations due to α₃DIV-$Zn^{2+}$ interactions is different from that of $Cd^{2+}$. Treatment with equimolar amount of EDTA recovers the spectrum of $Zn^{2+}$-free protein.*
